# Transcriptional regulation of genes involved in Zn transport after foliar Zn application to *Medicago sativa*

**DOI:** 10.1101/2020.05.11.088617

**Authors:** Alessio Cardini, Elisa Pellegrino, Philip J. White, Barbara Mazzolai, Marco C. Mascherpa, Laura Ercoli

## Abstract

Zinc (Zn) is an essential micronutrient for both plants and animals, and Zn deficiency is one of the most widespread problems for agricultural production. Although many studies have been performed on the biofortification of staple crops with Zn, few studies have focused on forage crops. In this study the molecular mechanisms of Zn transport-related in *Medicago sativa* L. were investigated following foliar Zn applications aimed at increasing the accumulation of Zn in edible tissues. Zinc uptake and redistribution between shoot and root were determined following the application of six Zn doses to leaves (0, 0.01, 0.1, 0.5, 1, 10 mg Zn plant^-1^). Twelve putative genes encoding proteins involved in Zn transport (*MsZIP_1-7_, MsZIF1, MsMTP1, MsYSL1, MsHMA4* and *MsNAS1*) were identified and the changes in their expression following foliar Zn application were quantified using newly designed RT-qPCR assays. Shoot and root Zn concentration was increased following foliar Zn applications ≥ 0.1 mg plant^-1^. Increased expression of *MsZIP2, MsHMA4* and *MsNAS1* in shoots, and of *MsZIP2* and *MsHMA4* in roots, was observed with the largest Zn dose. By contrast, *MsZIP3* was downregulated in shoots at Zn doses ≥ 0.1 mg plant^-1^. Three functional modules were identified in the *M. sativa* response to foliar Zn application: genes involved in Zn uptake by cells, genes involved in vacuolar Zn sequestration and genes involved in Zn redistribution within the plant. These results will inform genetic engineering strategies aimed at increasing the efficiency of crop Zn biofortification.

**One-sentence summary:** Upregulation of *ZIP2, NASI* and *HMA4* and downregulation of *ZIP3* are associated with Zn sequestration and shoot-to-root translocation in *Medicago sativa* following foliar Zn biofortification

## INTRODUCTION

Food quality is a key factor for human health (Geissler and Powers, 2017). For their well-being, humans require sufficient quantities of at least 18 mineral elements, which have specific physiological roles and are irreplaceable in the diet (White, 2016). Eight essential macronutrients (i.e., N, P, S, Ca, Mg, K, Na, Cl) are required in large amounts in the diet (> 100 mg day^-1^), and 10 micronutrients are required in smaller amount (e.g., Zn, Fe, F, Mn). The main sources of these elements in the human diet include edible crops, animal products (e.g., meat, fish, eggs) and dairy products (e.g., milk, cheese, butter) as well as mineral supplements (Keen, 1990; Prasad, 2013; White, 2016). A large proportion of the world’s population suffers from Zn related diseases (i.e., malabsorption syndrome, liver disease, chronic renal disease, sickle cell disease and other chronic diseases), since it relies on cereal-based diets with low Zn content due to poor soil Zn availability (WHO, 2005; Alloway, 2009; Prasad, 2013; Kumssa et al., 2015; Cakmak et al., 2017). Diversification of the human diet and biofortification of edible crops are therefore needed to alleviate Zn deficiency in humans. Similarly, increasing Zn concentrations in forage crops are important for maintaining livestock health and the quality of food products, which affect human health indirectly (McDonald et al., 2002; Ciccolini et al., 2017; Capstaff and Miller, 2018; Huma et al., 2019).

Zinc plays a major role as a co-factor of over 300 enzymes in plants and is an essential micronutrient (Broadley et al., 2007). Zinc is involved in various physiological functions, such as CO2 fixation, protein synthesis, free radical capture, regulation of growth and development, and disease resistance (Sasaki et al., 1998; Broadley et al., 2007). Many structural motifs in transcriptional regulatory proteins are stabilized by Zn, such as Zn finger domains (Albert et al., 1998). Zinc deficiency reduces crop production, as does Zn excess (White and Pongrac, 2017). Excessive Zn^2+^ can compete with other cations in binding to enzymes and for transport across membranes, thereby impairing cellular activities (White and Pongrac, 2017). Thus, the uptake of Zn^2+^ by cells and its transport within the plant must be strictly regulated. Plant cells have evolved several homeostatic mechanisms for avoiding Zn^2+^ toxicity when exposed to large Zn availability in their environment. These include the reduction of Zn influx to cells, the stimulation of Zn efflux from the cytosol, the sequestration of Zn in vacuoles, and the chelation of Zn by Zn binding ligands. In general, the concentration of Zn in plant tissues must be kept between 15 to 300 μg Zn g^-1^ dry matter (DM) to maintain cell structure and function (Broadley et al., 2012; White and Pongrac, 2017). Although tolerance to large tissue Zn concentrations varies among species (Alloway, 2008; White and Pongrac, 2017), Zn concentrations above 400-500 μg g^-1^ DM often cause toxicity symptoms including impaired root and shoot growth, chlorosis and necrosis of leaves, reduced photosynthesis, nutrient imbalance and ultimately loss of yield (Chaney, 1993; Broadley et al., 2007; Di Baccio et al., 2009; White and Pongrac, 2017).

The process of producing crops with greater mineral concentrations in edible tissues is called biofortification and provides a solution to the problem of mineral deficiencies in human and animal nutrition (White and Broadley, 2005). There are various approaches to Zn biofortification of edible crops, including agronomic strategies and conventional or transgenic breeding strategies. Agronomic biofortification aims to increase Zn concentrations in edible tissues through the application of Zn-fertilisers to the soil or to leaves. It is relatively inexpensive and efficient (Saltzman et al., 2013). Foliar application of Zn is generally more effective than the application of Zn fertilisers to soil, since Zn uptake by plant roots is often limited by the low solubility of Zn salts, its binding to organic substrates, and its immobilization in the microbial biomass (Gregory et al., 2017). Both agronomic and genetic biofortification strategies have been studied extensively in cereal staple crops, such as rice, wheat and maize, but less in legumes, such as beans, peas or lentils (White and Broadley, 2005, 2011; Rawat et al., 2013). An international program, the HarvestPlus Zinc Fertilizer Project, is exploring the potential of Zn fertilisers to enhance the yields and Zn concentrations in edible portions of staple crops in developing countries of Africa, Asia and South America (www.harvestzinc.org) (Cakmak, 2012), but this program does not include forage crops.

The natural direction of Zn flux in plants is from the soil via roots to the shoot and seeds (White and Broadley, 2009). Various transport proteins and ligands that are responsible for Zn^2+^ uptake by roots and its transport and sequestration within the plant have been characterized (Olsen and Palmgreen, 2014; Caldelas and Weiss, 2017; White and Pongrac, 2017). Among these, ZRT-IRT-like Proteins (ZIPs), have been studied in several plants, including *Arabidopsis thaliana*, soybean (*Glycine max*), barley (*Hordeum vulgare*) barrel medic (*Medicago truncatula*) and rice (*Oryza sativa*) (Grotz et al., 1998; Zhao and Eide, 1996; López-Millán et al., 2004; Milner et al., 2013; Tiong et al., 2015). These proteins not only transport Zn^2+^ across membranes, but can also transport other transition metal cations, including Cd^2+^, Fe^3+^/Fe^2+^, Mn^2+^, Ni^2+^, Co^2+^ and Cu^2+^ (Grotz et al., 1998; Mäser et al., 2001, Eckhardt et al., 2001). Generally, the expression of ZIP genes is upregulated when plants become Zn deficient (Ramesh et al., 2003; Ishimaru et al., 2006; Eide et al., 1996), facilitating Zn influx to cells and movement of Zn between organs, and also when plants become Fe or Mn deficient (Bughio et al., 2002; Vert et al., 2002; Ishimaru et al., 2006; Pedas et al., 2008). Other proteins that transport Zn include the Metal Tolerance Proteins (MTPs), which function as cation/proton antiporters and are thought to transport Zn into vacuoles (Kolaj-Robin et al., 2015) and the Yellow Stripe-Like Proteins (YSLs), which transport the Zn-Nicotianamine complex (NA-Zn) and load Zn into the xylem and phloem (Curie et al., 2009). The Zinc Induced Facilitators (ZIFs) and the Heavy Metal transporters (HMAs) are implicated in Zn influx to vacuoles and to the xylem, respectively (Olsen and Palmgren, 2014). Zinc is chelated by organic molecules, such as the carboxylic acid, citric acid, and nicotianamine (NA) in plants (Sinclair and Krämer, 2012). Nicotianamine is a non-proteinogenic amino acid with a high affinity for Fe, Cu and Zn, and is involved in their homeostasis (Deinlein et al., 2012). Nicotianamine mediates the intercellular and interorgan movement of Zn and was found to enable Zn hyperaccumulation in *Arabidopsis halleri* and *Noccaea caerulescens* (Deinlein et al., 2012; Foroughi et al., 2014) In general the functions of these transporters have been studied by expressing them in yeast, but to understand how the various Zn transport proteins and chelates act together to maintain appropriate cytosolic and tissue Zn concentrations it is important to study the responses of an intact plant to fluctuations in Zn supply.

In this study the transcriptional responses of genes encoding Zn transport-related processes facilitating Zn uptake by cells, vacuolar sequestration and redistribution within the plant were studied following foliar Zn application to the most productive and widely cultivated forage legume, alfalfa (*Medicago sativa* L.). The study was designed to provide information on the molecular responses to Zn biofortification of forage crops (Foyer et al., 2016; Capstaff and Miller, 2018). The following hypotheses were tested: i) foliar application of Zn increases shoot and root Zn concentrations, which results in changes in the expression of genes involved in Zn transport-related processes to detoxify excess Zn; ii) genes encoding Zn transport-related processes are organized in functional modules, that act in a concerted manner to redistribute Zn within the plant to maintain non-toxic cytosolic and tissue Zn concentrations. Genes encoding putative Zn transport-related processes were identified in alfalfa through phylogenetic comparisons and their likely roles are discussed. Changes in the expression of these genes following foliar Zn application were determined and the possible effects of these on the redistribution of Zn within cells and between tissues are discussed. The knowledge gained from this study could help to optimize Zn biofortification strategies when using foliar Zn fertilisers and to provide strategies for breeding forage crops to addresses Zn deficiencies in livestock.

## RESULTS

### Shoot and root Zn concentrations

The application of Zn to leaves did not modify shoot or root biomass and all *M. sativa* plants had root nodules (data not shown). However, Zn concentrations in both shoots and roots were strongly affected by foliar Zn application (F_(5,17)_=32.61, *P*<0.001; F_(5, 17)_=28.53, *P*<0.001; respectively) (Fig. 1). A foliar Zn application of 0.01 mg Zn plant^-1^ produced a shoot Zn concentration similar to that of the control (no-Zn addition), but shoot Zn concentrations were increased progressively by larger doses (0.1 < 0.5/1 < 10 mg Zn plant^-1^), from more than threefold to 35-fold more than that of the control (Fig. 1). Foliar applications of 0.01, 0.1 and 0.5 mg Zn plant^-1^ did not produce root Zn concentrations greater than that of the control treatment, but foliar doses of 1 and 10 mg Zn plant^-1^ increased root Zn concentrations to threefold and 11-fold more than the control treatment, respectively. Shoot and root Zn contents were also strongly affected by foliar Zn application (F_(5,17)_=53.73, *P*<0.001; F_(5, 17)_=32.45, *P*<0.001; respectively) and their responses to increasing foliar Zn applications followed the corresponding Zn concentrations (Supplemental Fig. S1).

**Figure 1.**
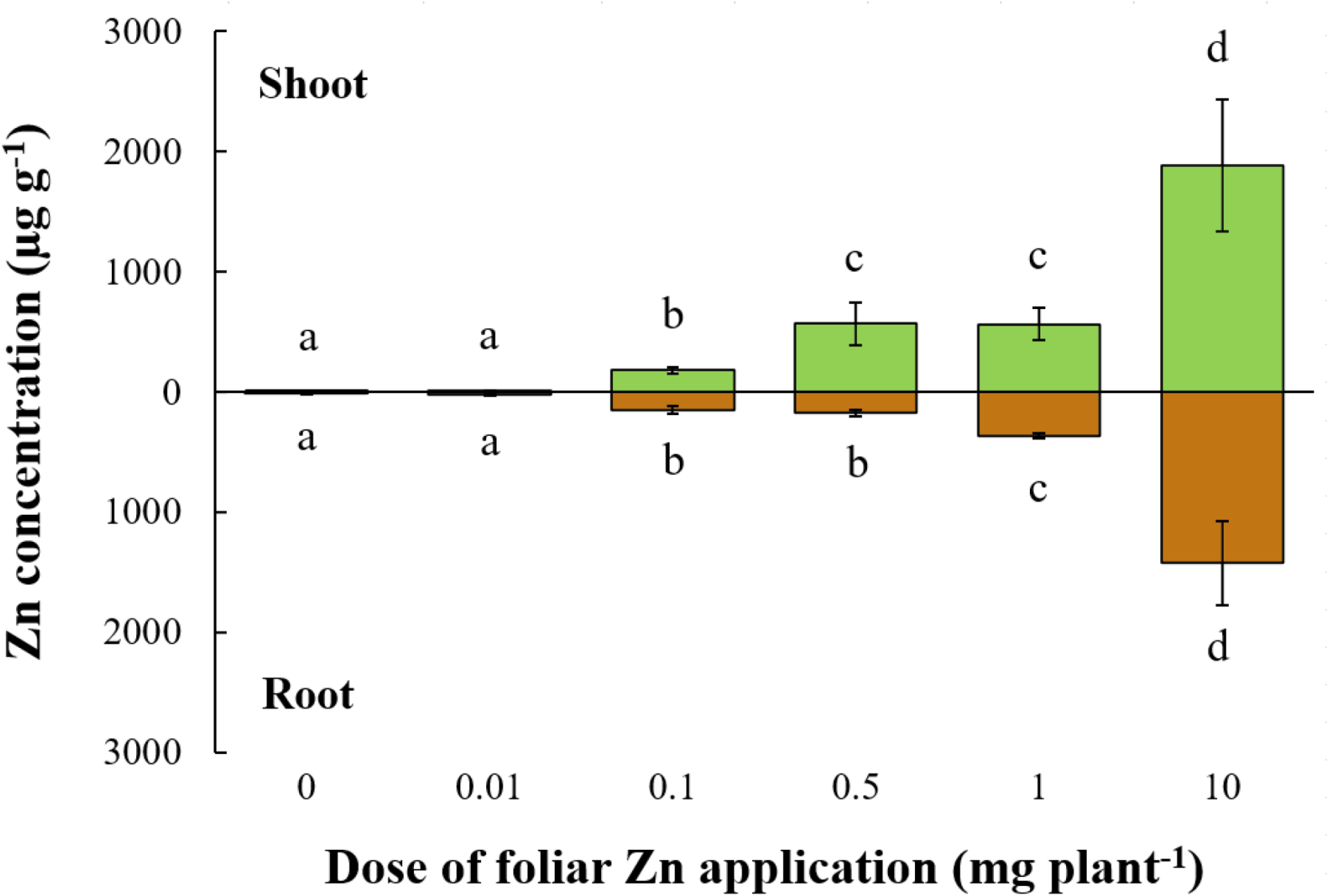
Zinc (Zn) concentrations in shoots and roots of alfalfa (*Medicago sativa*) five days after the application of Zn doses of 0, 0.01, 0.1, 0.5, 1 or 10 mg Zn plant^-1^ to leaves. Means ± standard error of three replicates are shown. Differences among the applied Zn doses were tested separately for shoot and root by one-way analysis of variance. Different letters denote significant differences in Zn concentrations in shoots and roots independently, according to Tukey-B honestly test (*P* < 0.05).

### Phylogenetic analysis

Phylogenetic analysis of the coding sequences of the *ZIP* genes revealed several distinct clades (Supplemental Fig S2). One clade contained sequences for *MsZIP2* and *MsZIP7,* which were similar to each other. In addition, the sequence of *MsZIP2* was closely related to those of *MtZIP2* and *GmZIP1-ZIP2* and the sequence of *MsZIP7* was closely related to those of *MtZIP7* and *AtZIP11*. Another clade contained the sequences of *MsZIP1, MsZIP3, MsZIP5* and *MsZIP6*. The sequence of *MsZIP1* clustered with that of *MtZIP1*. Sequences of *MsZIP3* and *MsZIP5* were similar to each other and clustered with the corresponding sequences for *M. truncatula* genes (Supplemental Fig. S2). Sequences for *MsZIP1, MsZIP3* and *MsZIP5* were closely related to each other, whereas that of *MsZIP6* formed a separate cluster with the sequences of *MtZIP6* and *AtZIP12*. The sequence of *MsZIP4* was distant from the sequences of other *M. sativa ZIPs* and formed a cluster with the sequences of *MtZIP4* and *AtZIP4*.

Phylogenetic analyses of the coding sequences of the other genes related to Zn transport processes revealed that they were all similar to their *M. truncatula* counterparts. As regards *ZIF*, the sequence of *MsZIF1* clustered with the sequences of *MtZIF1* and *GmZIF1* (Supplemental Fig. S3a). As regards *MTP*, the sequence of *MsMTP1* formed a cluster with *MtMTP1* and *GmMTP1* and was also related to *AtMTP1* and *AtMTPA1* (Supplemental Fig. S3b). Similarly, the sequence of *MsYSL1* was most similar to those of *MtYSL1* and *GmYSL1* (Supplemental Fig. S3c) and the sequence of *MsHMA4* was most similar to those of *MtHMA4* and *GmHMA4* (Supplemental Fig. S3d). Finally, the sequence of *MsNAS1* was closely related to those of *MtNAS* and *GmNAS* (Supplemental Fig. S3e).

### Gene expression analysis

The expression of *MsZIP3* was significantly downregulated at foliar doses of 0.1, 1 and 10 mg Zn plant^-1^ (F_(3, 11)_ = 28.46, *P*<0.01) (Fig. 2). By contrast, the expression of *MsZIP2* was significantly upregulated at the largest dose of 10 mg Zn plant^-1^ (F_(3, 11)_ = 5.59, *P*<0.05). The expression of *MtZIP1, MtZIP5* and *MtZIP6* in shoots was not significantly affected by foliar Zn application, although a general trend towards downregulation with increasing foliar Zn doses was observed. The expression of *MsZIP4* and *MsZIP7* in shoots was unaffected by foliar Zn application. The expression of *MsZIP2* was significantly upregulated in roots at the largest foliar dose of 10 mg Zn plant^-1^ (F_(3, 11)_=9.26, *P*<0.01 (Fig. 2). In roots, *ZIP* genes were not significantly affected by foliar Zn application, although a general trend of *MtZIP1, MtZIP3, MtZIP5* and *MtZIP7* towards upregulation with increasing foliar Zn doses was observed. Of the other genes related to Zn transport processes, the expression of *MsHMA4* was significantly upregulated in both shoots (F_(3, 11)_ = 115.29, *P*<0.01) and roots (F_(3, 11)_ = 14.23, *P*<0.01) following the application of 1 and 10 mg Zn plant^-1^ (shoots: +63% and +424%, respectively; roots: +86% and +66%, respectively; Fig. 3). In shoots, the expression of *MsHMA4* was about 3-fold higher following a dose of 10 mg Zn plant^-1^ than following a dose of 1 mg Zn plant^-1^, whereas the expression of *MsHMA4* in roots was similar when 1 or 10 mg Zn plant^-1^ was applied. The expression of *MsNAS1* was also significantly upregulated (F_(3, 11)_ = 6.46, *P* < 0.05) at the largest foliar Zn dose (10 mg plant^-1^), whereas its expression in roots was unaltered following foliar Zn application (Fig. 3). In shoots, *MsYSL1* and *MsZIF1* were not significantly affected by foliar Zn application, although there was a trend towards upregulation of the expression with increasing Zn doses, while the expression of *MsMTP1* remained unaltered following the application of Zn (Fig. 3). Finally, in roots, *MsMTP1* and *MsZIF1* were not significantly affected by foliar Zn application, although there was a trend towards upregulation of the expression with increasing Zn doses, whereas the expression of *MsYSL1* remained unchanged (Fig. 3).

**Figure 2.**
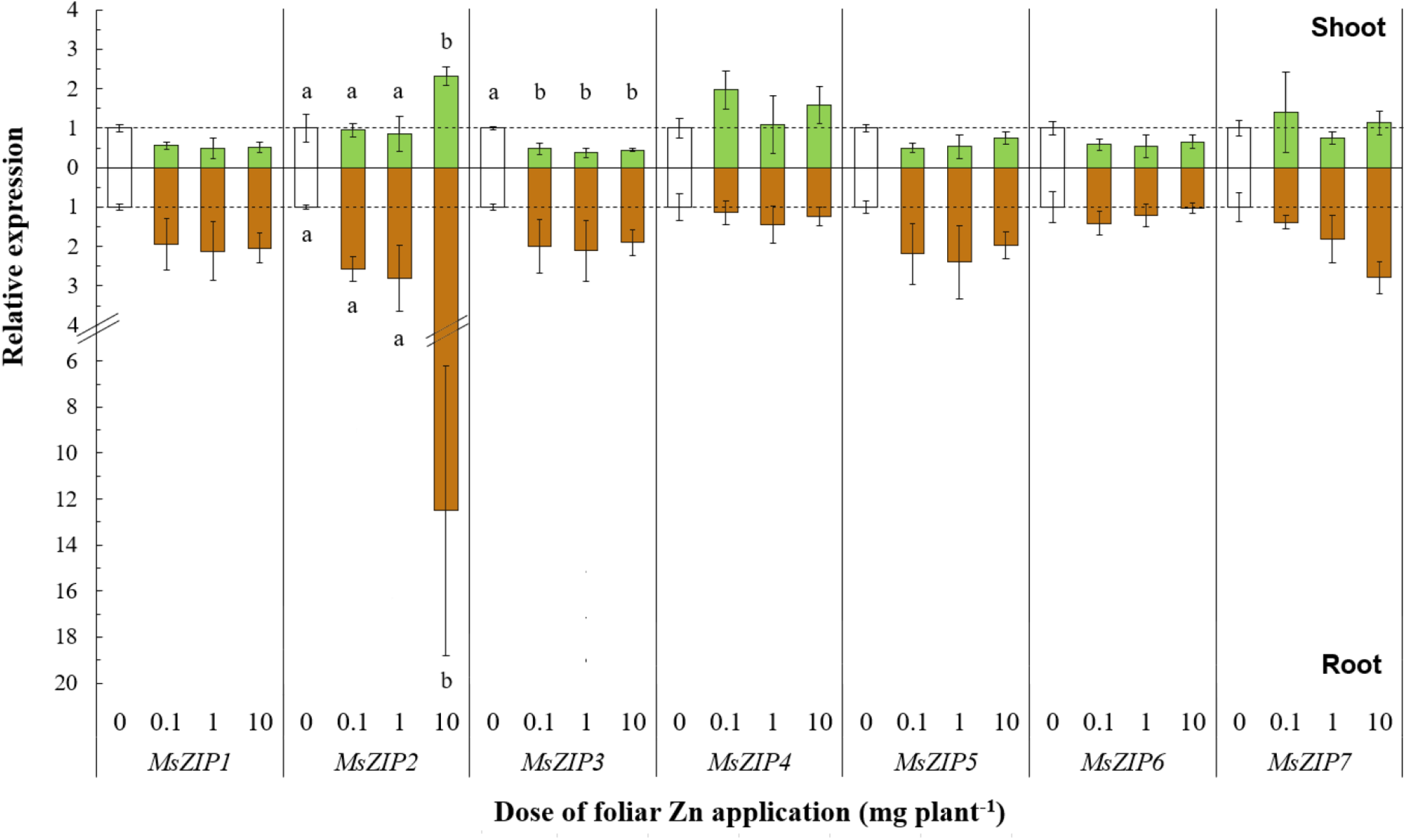
Relative expression of seven transmembrane zinc (Zn) transporter genes (*MsZIP_1-7_*) five days after the application of Zn doses of 0, 0.1, 1 or10 mg Zn plant^-1^ to leaves of alfalfa (*Medicago sativa*). Means ± standard error of three replicates are shown. The expression levels were calculated relative to reference genes (*MsACT-101* for shoot and *MsEF1-α* for root) and to the control (0 mg Zn plant^-1^). The broken line denotes the threshold between up- and down-regulation relative to the control. Differences in the expressions of each gene after different Zn doses were tested separately for shoot and root by one-way analysis of variance. Different letters denote significant differences among Zn doses, according to Tukey-B test (*P* < 0.05). The full names of the genes are reported in the legend of Figure 1.

**Figure 3.**
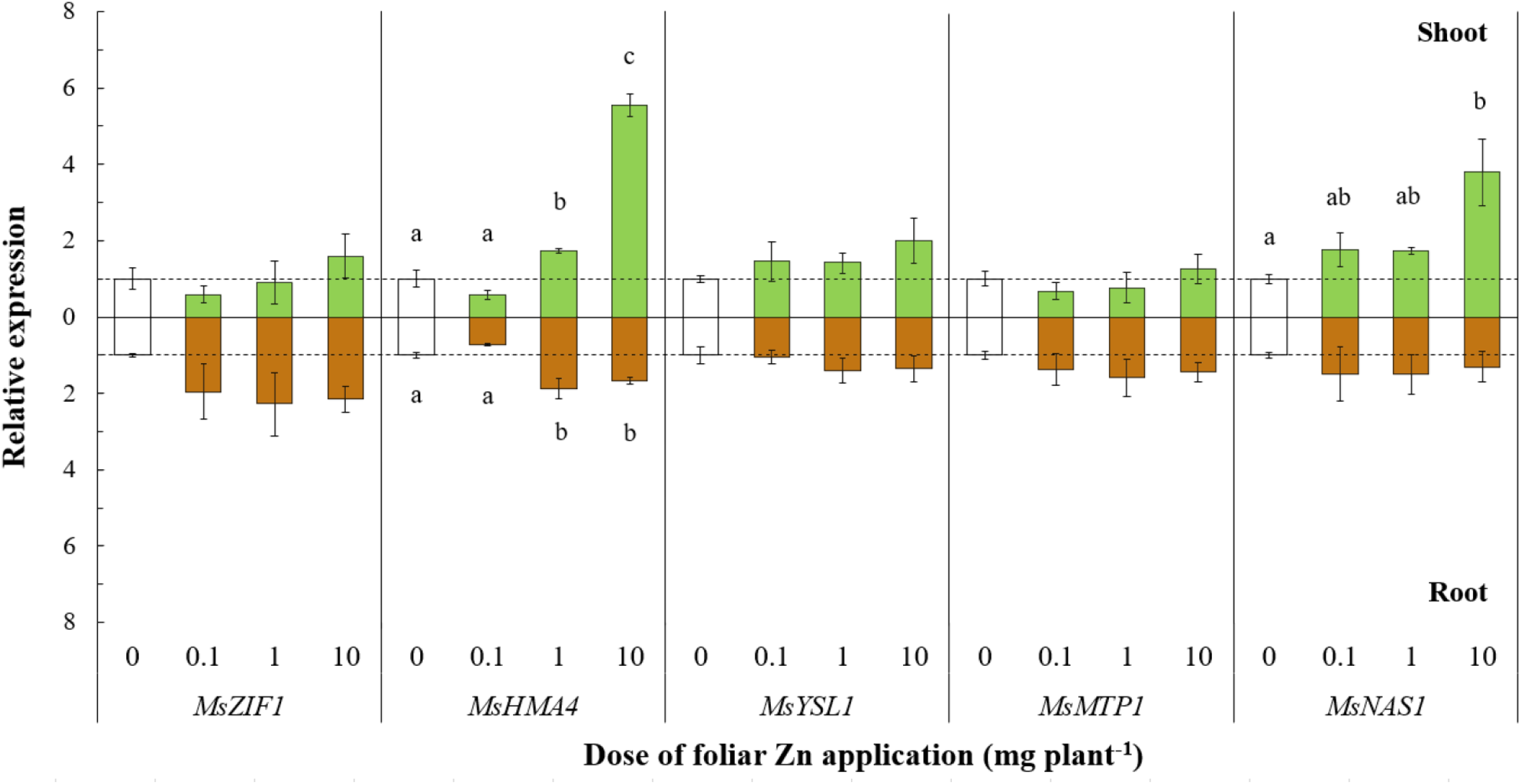
Relative expression of genes related to Zn transport processes (*MsZIF1*, *MsHMA4*, *MsYSL1, MsMTP1* and *MsNAS1*) five days after the application of Zn doses of 0, 0.1, 1 and 10 mg Zn plant^-1^ to leaves of alfalfa (*Medicago sativa*). Means ± standard error of three replicates are shown. The expression levels were calculated relative to reference genes (*MsACT-101* for shoot and *MsEF1-α* for root) and to the control (0 mg Zn plant^-1^). The broken line denotes the threshold between up- and down-regulation relative to the control. Differences in the expression of each gene at the different Zn doses were tested separately for shoot and root by one-way analysis of variance. Different letters denote significant differences among Zn doses, according to Tukey-B test (*P* < 0.05). The full names of the genes are reported in the legend of Figure 1.

Using correlation analysis to reveal functional modules of genes whose expression is coregulated in plants, three functional modules for ZIP gene co-expression were observed (Fig. 4a; *r* > 0.6). In the first functional module, the expression of *MsZIP1, MsZIP3, MsZIP5* and *MsZIP6* in shoots were all strongly correlated. Although the expression of *MsZIP7* in shoots was also placed in this module, its expression was poorly correlated with the expression of other *ZIP* genes in either shoots or roots. The second functional module comprised correlations between the expression of *MsZIP4* in shoots and roots, and the expression of *MsZIP1*, *MsZIP3*, *MsZIP4*, *MsZIP5* and *MsZIP6* in roots. The third functional module comprised correlations in the expression of *MsZIP2* in shoots and roots and *MsZIP7* in roots. With respect to the expression of the other genes involved in Zn transport related processes, there was a strong divergence in their expression patterns in shoots and roots (Fig. 4b). In the root, the expression of *MsNAS1*, *MsYSL1, MsZIF1* and *MsMTP1* were strongly correlated (Fig. 4b). The expression of *MsYSL1*, *MsZIF1* and *MsMTP1* in roots showed good correlations with the expression of *MsMTP1* and *MsZIF1* in shoots. In the shoot the expression of *MsYSL1*, *MsMTP1* and *MsZIF1* were all strongly correlated. Also in shoots, the expression of *MsNAS1* and *MsHMA4* were well correlated and showed a good correlation with the expression of *MsMTP1* and *MsZIF1* in shoots. Interestingly, the expression of *MsNAS1* strongly correlated with that of *MsYSL1* in shoots.

**Figure 4.**
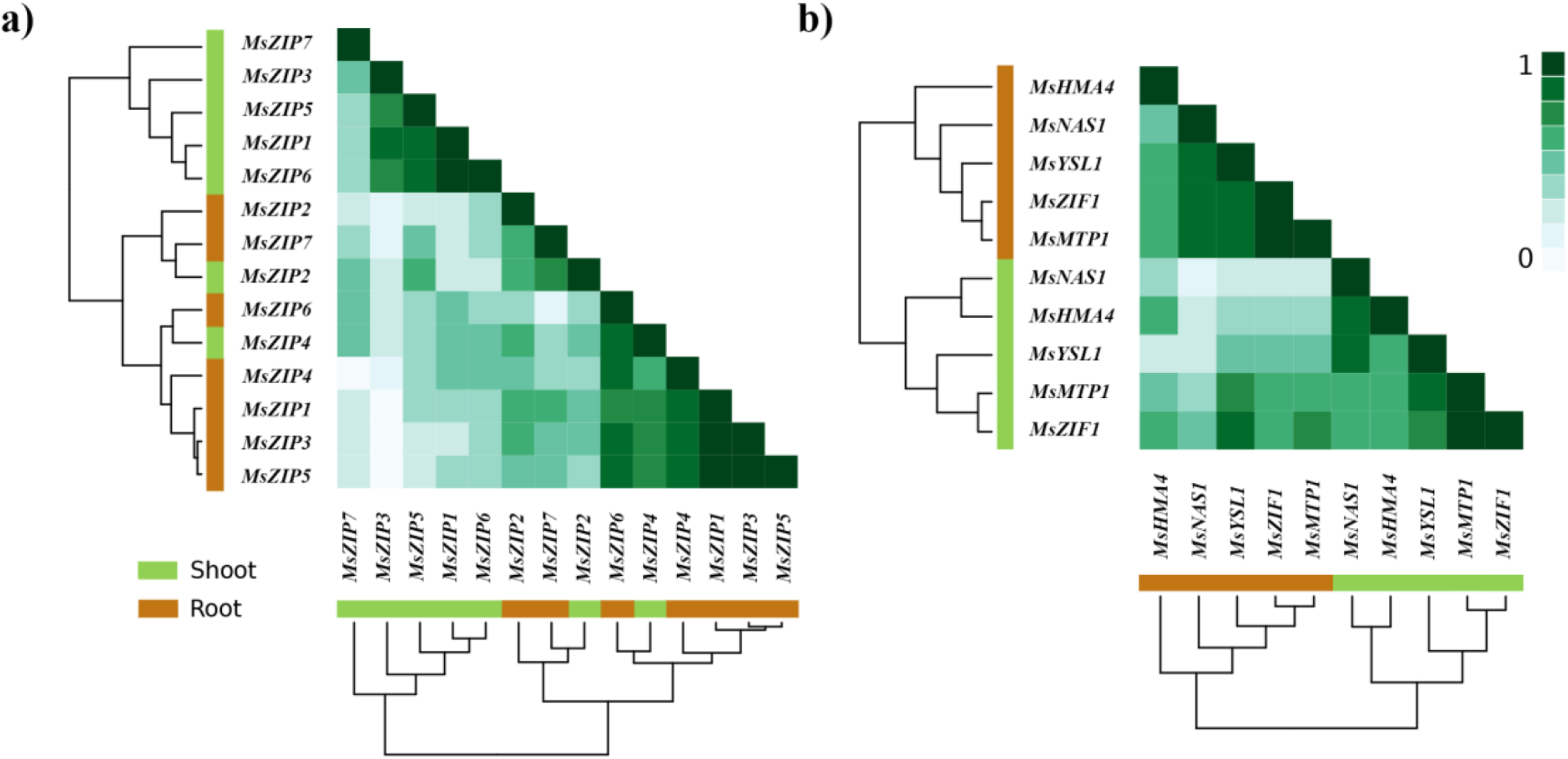
Heatmaps reporting the correlations between the differences in expression of genes related to zinc (Zn) transport processes in alfalfa (*Medicago sativa*) after foliar application of Zn doses of 0.1, 1 and 10 mg Zn plant^-1^ relative to a control dose of 0 mg Zn plant^-1^. The similarity in the degree of correlation in fold-change of gene expression to Zn application relative to the control is based on the average linkage clustering of the Pearson correlations (r). In the clustering trees the genes are indicated in brown for roots and in green for shoots, while the ranks of correlations of the heatmap are indicated by color intensity (r 0 to 1: from low to strong intensity of green). Seven genes encoding transmembrane Zn transporter (*MsZIP_1-7_*) (a); four genes encoding cellular Zn transporters (including vacuolar transporters) (*MsZIF1*, *MsHMA4*, *MsYSL1* and *MsMTP1*) and a gene encoding a nicotianamine synthase (*MsNAS1*) (b).

PERMANOVA showed that the expression of *ZIP* genes was significantly affected by foliar Zn application dose and differed between shoots and roots, which explained 29% and 23% of the total variance, respectively (Table 1). The expression of other genes related to Zn transport processes that were studied (*MsZIF1, MsNAS1, MsHMA4, MsYSL1* and *MsMTP1*) were also affected by foliar Zn application dose and the organ examined. Zinc application dose explained 17% of the total variance, while plant organ explained 19%. PERMANOVA on all studied genes highlighted a significant effect of Zn application dose, plant organ and their interaction on gene expression, explaining 68% of the total variance.

**Table 1.** Permutation analyses of variance (PERMANOVAs) on the effect of application of three doses of zinc (Zn) (0.1, 1 and 10 mg Zn plant^-1^) and plant compartment (shoot and root) on the expression of seven *MsZIP* genes and separately on the expression of other five genes (*MsZIF1, MsNAS1, MsHMA4, MsYSL1* and *MsMTP1*) (see Table 1, Fig. 1). A PERMANOVA was also performed on the response of all the genes. The analysis of homogeneity of multivariate dispersion (PERMDISP) was also performed. The studied plant was alfalfa (*Medicago sativa* L.). The analysis included also no-Zn addition control. Gene relative expression was studied on a total of 24 experimental units, corresponding to three replicates per each level of treatment.

| Response variables | Explanatory variables | Zn application (Zn) | Plant compartment (Comp) | Zn x Comp | Residual |
| --- | --- | --- | --- | --- | --- |
| *ZIP genes* | Pseudo F | 5.56 | 8.16 | 1.76 |  |
|  | *P*(perm) | **0.002** | **0.001** | 0.082 |  |
|  | Explained variance (%) | 29.1 | 22.9 | 9.7 | 38.3 |
|  | PERMDISP *P*(perm) | **0.005** | 0.766 |  |  |
| Other genes | Pseudo F | 3.06 | 5.59 | 1.76 |  |
|  | *P*(perm) | **0.007** | **0.015** | 0.1 |  |
|  | Explained variance (%) | 17.3 | 19.35 | 12.78 | 50.55 |
|  | PERMDISP *P*(perm) | 0.412 | 0.852 |  |  |
| All genes | Pseudo F | 4.27 | 10.49 | 3.41 |  |
|  | *P*(perm) | **0.001** | **0.001** | **0.003** |  |
|  | Explained variance (%) | 17.3 | 25.2 | 25.6 | 31.9 |
|  | PERMDISP *P*(perm) | 0.152 | **0.030** |  |  |

## DISCUSSION

### Plant Zn nutritional status after foliar Zn application

The critical leaf concentration for Zn deficiency approximates 15–20 μg Zn g^-1^ dry weight and the critical leaf concentration for Zn toxicity approximates 400-500 μg Zn g^-1^ (Broadley et al., 2012; White and Pongrac, 2017). Before foliar Zn application, the alfalfa plants used in the experiments reported here were probably Zn deficient, since their shoot Zn concentrations were below the critical leaf concentration for Zn deficiency (Fig. 1). After the application of the lowest foliar Zn dose (0.01 mg plant^-1^) plants probably remained Zn deficient (7.6 μg Zn g^-1^ dry weight), but all other foliar Zn doses increased Zn concentrations in shoots above the critical concentration for Zn deficiency (Fig. 1). Plants treated with 0.1 mg Zn plant^-1^ probably had an optimal Zn status for plant growth, whereas plants treated with 0.5 and 1 mg Zn plant^-1^ had shoot Zn concentrations close to the toxicity threshold. When a foliar dose of 10 mg Zn plant^-1^ was applied, shoot Zn concentrations greatly exceeding the threshold for Zn toxicity (Fig. 1). Plants often exhibit characteristic visual symptoms of Zn deficiency and Zn toxicity when these occur (Broadley et al., 2012; White and Pongrac, 2017), but five days after foliar Zn application no visual symptoms of Zn deficiency or toxicity, nor differences in plant biomass, were observed among plants receiving contrasting foliar Zn doses (data not shown). Foliar Zn doses larger than 0.1 mg Zn plant^-1^ resulted in incremental increases in the Zn concentration of roots (Fig. 1), despite Zn having limited mobility in the phloem (White and Broadley, 2011; White, 2012). This observation suggests that roots can act as a sink for Zn applied to leaves, thereby mitigating excessive Zn accumulation in shoot tissues. In previous work, foliar application of Zn was shown to increase Zn concentration in phloem-fed tissues, such as fruits, seed, and tubers (Cakmak, 2004, 2008; Cakmak et al., 2010; White et al., 2017). The shoot to root Zn concentration ratio shifted from values below one in conditions of Zn deficiency (0.4) to values greater than one in Zn-replete or Zn-intoxicated plants (1.3 – 3.2) (Fig. 1). When the plants are Zn deficient the recirculation of Zn between organs via the xylem and phloem is required to meet minimal growth demands and the application of foliar Zn to Zn deficient plants must be effectively redistributed within the plant (Erenoglou et al., 2011; Sinclair and Kramer, 2012), whereas when excessive foliar Zn is applied, Zn must be chelated in the cytoplasm, sequestered in the vacuole and redistributed via the phloem or xylem to other organs to avoid toxicity (White and Pongrac, 2017).

### Foliar Zn application alters the expression of genes involved in Zn transport-related processes

Despite several genes encoding Zn transporters having been identified in plants, and the encoded proteins characterized, the mechanisms of Zn uptake and transport in alfalfa are still largely unknown. However, the recently sequenced alfalfa genome has allowed the discovery of genes involved in Zn uptake and distribution within this species (O’Rourke et al., 2015). In the present study, 12 putative genes encoding proteins likely to be involved in processes related to Zn transport in the plant (*MsZIP_1-7_*, *MsZIP1*, *MsMTP1*, *MsYSL1*, *MsHMA4* and *MsNAS1*) were identified and their expression in shoots and roots quantified following foliar Zn applications (Fig. 2; Fig. 3). The expression of genes encoding proteins involved in Zn uptake by cells, vacuolar sequestration and redistribution within the plant responded differently to increasing foliar Zn dose. The expression of *MsZIP2*, *MsHMA4* and *MsNAS1* in shoots was increased by increasing foliar Zn dose, while only the expression of *MsZIP2* and *MsHMA4* in roots were upregulated by increasing foliar Zn dose (Fig. 2; Fig. 3). By contrast, *MsZIP3* was downregulated in shoots when foliar Zn doses ≥ 0.1 mg Zn plant^-1^ were applied (Fig. 2). These changes in gene expression might produce a reduction in Zn uptake capacity (but not necessarily a reduction in actual Zn uptake since this will also be related to apoplastic Zn concentration) by cells in both the shoot and root (by reducing Zn influx through *MsZIP3* and increasing Zn efflux through *MsHMA4*), chelation of Zn using Zn-NA in the shoot for Zn-detoxification and phloem transport, and greater recirculation of Zn within the plant in both the phloem and xylem (by increasing NA concentrations and *MsZIP2* and *MsHMA4* activities), and are, therefore, consistent with the observation that increasing foliar Zn dose increases both shoot and root Zn concentrations.

### Regulation of genes encoding ZIP transporters

The influx and efflux of Zn across the plasma membrane of plant cells must be tightly controlled to allow optimal cell functioning and hence to ensure normal plant growth and development (Sinclair and Krämer, 2012). The expression of only two of the seven *ZIP* genes studied, *MsZIP2* and *MsZIP3*, showed statistically significant responses to foliar Zn application (Fig. 2). The expression of *MsZIP2* was significantly upregulated in both shoots and roots in response to the largest dose of foliar Zn applied (10 mg Zn plant^-1^). It is likely that this dose is toxic to both shoot and root cells. The relative induction in the expression of *MsZIP2* was greatest in roots. The phylogenetic analysis of ZIP transporters revealed that *MsZIP2* is closely related to *MtZIP2* and *AtZIP2* (Supplemental Fig. S6). Thus, *MsZIP2* is probably located in the plasma membrane performing similar functions to *MtZIP2* and *AtZIP2*. Burleigh et al. (2003) reported that *M. truncatula* plants grown with adequate soil Zn availability expressed *MtZIP2* in roots and stems, but not in leaves. The expression of *MtZIP2* in roots increased with increasing Zn fertiliser applications to soil, with the greatest expression being found at toxic Zn doses (Burleigh et al., 2003). Similarly, Milner et al. (2013) found that the expression of *AtZIP2* was ~10-fold higher in roots than shoots in Zn-replete *Arabidopsis thaliana* plants and that Zn deficiency reduced the expression of *AtZIP2* in both roots and shoots. The localization of *ZIP2* at the plasma membrane was observed in both *M. truncatula* (Burleigh et al., 2003) and *A. thaliana* (Milner et al., 2013). The expression of *AtZIP2* was localized to the stele of the root (Milner et al., 2013), supporting a role of *AtZIP2* in long distance transport of Zn between roots and shoots. It is possible that the increased expression of *MsZIP2* observed in our study when plants experience Zn toxicity might be a detoxification strategy, either through storing excess Zn in xylem parenchyma cells or recirculating Zn in the xylem.

The expression of *MsZIP3* was significantly downregulated in shoots following the foliar application of Zn (Fig. 2). The ZIP3 transporter is thought to mediate Zn influx to the cell from the apoplast (Sinclair and Kramer, 2012). Therefore, the downregulation of *MsZIP3* in shoots of plants receiving more Zn is consistent with the ability of plant cells to control their Zn uptake to effect cytoplasmic Zn homeostasis. Reduced expression of *MsZIP3* in plants with a greater Zn supply is also in agreement with previous studies of *M. truncatula* and *A. thaliana* (Grotz et al., 1998; Lopéz-Millán et al., 2004). However, although *AtZIP3* could restore growth to a Zn-uptake defective yeast (Milner et al., 2013), *MtZIP3* was not found to be able to restore the growth of a Zn-uptake defective yeast in Zn-limited media, although it did restore the growth of a Fe-uptake defective yeast in Fe-limited media (López-Millán et al., 2004). Thus, the MsZIP3 transporter could have a higher affinity for Fe than Zn. In *O. sativa ZIP3* gene is expressed in the xylem parenchyma and transfer cells and might be responsible for unloading transition metal cations from the xylem to the parenchyma in plants receiving an excessive Zn supply (Sasaki et al., 2015). The role of OsZIP3 in unloading Zn from the vascular tissues, suggests that the reduced expression of *MsZIP3* in shoots of *M. sativa* receiving an excessive foliar Zn dose might be a detoxification strategy to reduce Zn uptake by shoot cells.

The observation that foliar Zn applications had no effect on the expression of ZIP genes, except *MsZIP2* and *MsZIP3* (Fig. 2), might be explained by the roles of ZIP proteins in the transport of other transition metals. For example, evidence of Cu and Mn transport by *ZIP4* were provided through yeast complementation studies (Wintz et al., 2003; López-Millán et al., 2004). Moreover, applying the same technique, a role of *ZIP6* was highlighted in the transport of Fe by López-Millán et al. (2004), whereas Wintz et al. (2003) did not find any involvement of *ZIP6* in the transport of Cu, Zn or Fe. Although the changes in the expression of *MsZIP1*, *MsZIP5* and *MsZIP6* following foliar Zn application were not statistically significant, changes in their expression in shoots were positively correlated with changes in the expression of *MsZIP3,* showing a general trend for them to be downregulated following foliar Zn application and suggesting that these four ZIPs might act as a functional module in the shoot (Fig. 4). By contrast, the expression of *MsZIP1, MsZIP3, MSZIP4,* and *MsZIP5* were positively correlated in roots, suggesting that these genes behave as a functional module in roots (Fig. 4).

### Regulation of genes encoding other Zn transport-related processes

The expression of *MsHMA4,* which is implicated in Zn redistribution within the plant (Hussain et al., 2004; Sinclair et al., 2018), was increased in both shoots and roots of plants whose shoot Zn concentration suggested they were close to, or experiencing, Zn toxicity (Fig. 3). The significant upregulation of *MsHMA4* following foliar application of ≥ 1 mg Zn plant^-1^ might be related to the removal of excess Zn from both shoots and roots. This interpretation is consistent with the role of HMA4 in *A. thaliana* and in the metal hyperaccumulators *Arabidopsis halleri* and *Noccea caerulescens* (Baker and Whiting, 2002; Hussain et al., 2004; Hanikenne et al., 2008; Ò Lochlainn et al., 2011; White and Pongrac, 2017), in which greater expression of *HMA4* results in greater Zn flux to the xylem and Zn translocation to transpiring leaves. However, the phylogenetic similarity of *MsHMA4* to *MtHMA4* and, particularly, to *AtHMA5* (Supplemental Fig. S3d) suggest a role in Cu transport (Andrés-Colás et al., 2006; Sankaran et al., 2009; Hermand et al., 2014). This implication of the latter observation is unclear.

Since Zn^2+^ concentrations are low in the alkaline phloem sap, the transport of most Zn in the phloem is as Zn ligand complexes, such as zinc-nicotianamine (NA-Zn) (Deshpande et al., 2018). Nicotianamine is the main Zn chelate in phloem transport and is also important for Zn sequestration in vacuoles (Deinlein et al., 2012), and tolerance of excessive Zn uptake (Aarts et al., 2014). Nicotianamine concentrations generally correlate with those of *NAS* transcripts, and for this reason *NAS* expression can be used as a proxy for NA content (Talke et al., 2006; Haydon et al., 2012). Accordingly, in the work reported here the increased expression of *MsNAS1* in shoots following the application of ≥ 1 mg Zn plant^-1^ (Fig. 3) probably reflects the role of NA in Zn detoxification through its sequestration within vacuoles and its redistribution from shoot to root after excessive foliar Zn applications. This observation is consistent with Deshpande et al. (2018), who found that the expression of *NAS2* in the durum wheat (*Triticum durum* Desf.) increased following foliar Zn application and reports that *NAS* expression is constitutively high in plants that hyperaccumulate Zn (Becher et al., 2004; Weber et al., 2004; Haydon et al., 2012; White and Pongrac, 2017).

Homologs of *MsMTP1* and *MsZIF1* were previously found to encode transporters loading Zn and NA into the vacuoles of *Thlaspi geosingense* and *A. thaliana* cells, respectively (Gustin et al., 2009; Haydon et al., 2012). Unexpectedly, the expression of these genes was unaffected by foliar Zn application (Fig. 3). This observation suggests that the proteins encoded by these genes might not contribute to Zn detoxification in *M. sativa*. Nevertheless, only *MsZIF1* of all the genes studied here showed a trend towards increased expression in roots with increasing foliar Zn dose (Fig. 3), which might indicate a role in detoxification of excess Zn in roots through its sequestration with NA in the vacuole. The high correlation between the expression of *MsMTP1* and *MsZIF1* in both shoots and roots suggests that they might constitute a functional module and act synergistically to sequester Zn in the vacuole (Fig. 4), as supported by other studies (Gustin et al., 2009; Haydon et al., 2012; Sharma et al., 2016).

In *A. thaliana*, *AtYSL1* has a role in the long-distance transport of the NA-Zn complex and in loading Zn into seeds (Jean et al., 2005; Curie et al., 2009). For this reason, an increase in the expression of *MtYSL1* was expected to occur in parallel with the increased expression of *MsNAS1* in shoots. However, the expression of *MsYSL1* did not show any significant change in shoots or roots in response to foliar Zn application, although there was a trend towards greater *MsYSL1* expression in shoots with increasing foliar Zn doses (Fig. 3). In addition, the high correlation in the expression of *MsNAS1* and *MsYSL1* in shoots in response to foliar Zn applications (Fig. 4) supports the expectation that these genes are components of a functional module affecting the long-distance transport of Zn in the plant, as it was previously highlighted in *A. thaliana* by Pita-Barbosa et al. (2019).

The responses of gene expression to foliar Zn applications suggest three functional modules that effect cytoplasmic Zn homeostasis through Zn transport related processes in *M. sativa*: genes involved in Zn influx to cells (shoots: *MsZIP1, MsZIP5,* and *MsZIP6;* roots: *MsZIP1, MSZIP4, MsZIP5*, and *MsZIP6*), genes involved in Zn sequestration in the vacuole (shoots and roots: *MsMTP1* and *MsZIF1*) and genes involved in Zn redistribution within the plant (shoots and roots: *MsHMA4* and *MsYSL1*).

In conclusion, this is the first study to characterise the expression of genes related to Zn transport processes following foliar Zn application to a forage legume and provides new molecular insights to the responses of Zn transport related processes to foliar Zn applications. A significant increase in the expression of *MsZIP2* as foliar Zn doses increase suggests the detoxification of excess Zn through the accumulation of Zn in xylem parenchyma cells. A decrease in the expression of *MsZIP3* as foliar Zn doses increase suggests a reduction in the Zn influx capacity of shoot cells to reduce Zn uptake. An increase in the expression of *MsHMA4* in roots and shoots as foliar Zn doses increase suggests an increase in the transport of Zn in the xylem when plants are subject to Zn toxicity, while an increase in the expression of *MsNAS1* in the shoot suggests the chelation of excess Zn in the shoot, enabling Zn sequestration in vacuoles or the redistribution of Zn to roots via the phloem. The elucidation of three functional modules of genes involved in (a) Zn influx to cells, (b) sequestration of Zn in the vacuole and (c) redistribution of Zn within the plant are fundamental to understanding the molecular mechanisms of cytoplasmic Zn homeostasis and might inform the selection of appropriate genotypes enabling greater Zn accumulation in edible portions or increased tolerance of Zn in the environment.

## MATERIALS AND METHODS

### Plant growth and experimental design

Surface sterilized seeds of alfalfa (*M. sativa* L.) were germinated on moist sterilized silica sand (1-4 mm size) in a climatic chamber at 24/21 °C day/night temperature, 16/18 h light/dark cycle and 200 μmol photons m^-2^ s^-1^. After two weeks of growth, three seedlings were transplanted to 1500 mL volume pots, filled with sterilized silica sand (number of pots 18) and *Sinorhizobium meliloti* was supplied as a filtrate to all plants to ensure that the plants produced nodules in all treatments. A Hoagland nutrient solution lacking Zn (Li et al., 2013) was used to fertilize the plants, with 10 mL solution being applied every week. After two months of growth, plants were treated with one of six doses of Zn (0, 0.01, 0.1, 0.5, 1 and 10 mg Zn plant^-1^) (three replicates per dose). Six ZnSO_4_·7H_2_O solutions of 0, 0.05, 0.5, 2.5, 5, 50 g Zn L^-1^ were prepared to supply these doses. A drop of Tween 20 detergent was added to the six solutions to break the surface tension of the leaves and enhance Zn uptake. Zinc was applied to the middle leaf laminae of the three plants in each pot as twenty 10 μl-droplets. The experiment was arranged in a fully randomized design, with three replicates for each Zn dose. The shoots and roots of the plants were harvested separately five days after Zn application. At harvest, 1 mM CaCl_2_ solution and water were used to remove any residual Zn from the leaf surface (Yilmaz et al., 2017). Shoot and root fresh weight was measured, whereas shoot and root dry weight was determined on subsamples after oven drying at 70°C to constant weight.

### Measurement of zinc concentrations

Approximately 100 mg of shoot or root dry biomass were carefully weighed and mineralized in a microwave medium pressure digestor (Milestone Start D, FKV Srl, Torre Boldone, Italy) with 7 mL of 69% HNO_3_ and 2 mL of 30% H_2_O_2_ (ultrapure grade). Zinc concentration in the resulting solutions was determined by inductively coupled plasma optical emission spectroscopy (ICP-OES) using an Optima 8000 spectrometer (Perkin Elmer, Waltham, MA, USA), following the procedure of Nölte (2003).

### Gene selection and design and validation of new RT-qPCR assay

Seven genes encoding putative ZRT-IRT-like proteins (ZIP) were selected for investigation (i.e., *ZIP1-7*) (Fig. 5). The selection was based on information gathered by Burleigh et al. (2003) and López-Millán et al. (2004) on the expression of genes encoding Zn transporters in the model legume *Medicago truncatula* and on the structure of the neighbor joining (NJ) tree built using available ZIP sequences of several plant species. Five more genes, whose products are involved in Zn transport-related processes (Olsen and Palmgren, 2014), were also chosen for investigation based on information gathered by other authors and on sequence similarity with other plant species. The *NAS1* gene encoding nicotianamine synthase (NAS) was chosen because this enzyme synthesizes nicotianamine (NA), which is involved in long-distance Zn transport (Curie et al., 2009) (Fig. 5). The *HMA4* gene, which encodes a transmembrane P-type ATPase heavy metal transporter, was chosen because this transporter loads Zn into the xylem in roots for its transport to shoots (Palmer and Guerinot, 2009) (Fig. 5). The *MTP1* gene, which encodes a transporter of the CDF family, was selected because this transporter is implicated in the sequestration of excess Zn in the vacuole (Desbrosses-Fonrouge et al., 2005; Gustin et al., 2009) (Fig. 5). The *ZIF1* gene, which encodes the Zn-induced facilitator 1 transporter, was chosen because it transports NA into the vacuole to chelate vacuolar Zn (Haydon and Cobbett, 2007; Haydon et al., 2012) (Fig. 5). The *YSL1* gene, which encodes a transporter of Zn-NA complexes, was chosen because it is implicated in Zn loading and transport of Zn in the phloem (Palmer and Guerinot, 2009) (Fig. 5). To standardize the expression of genes encoding Zn transport-related processes, two reference genes were selected: actin (*ACT*) and elongation factor 1-α (*EF1-α*) (Nicot et al., 2005).

**Figure 5.**
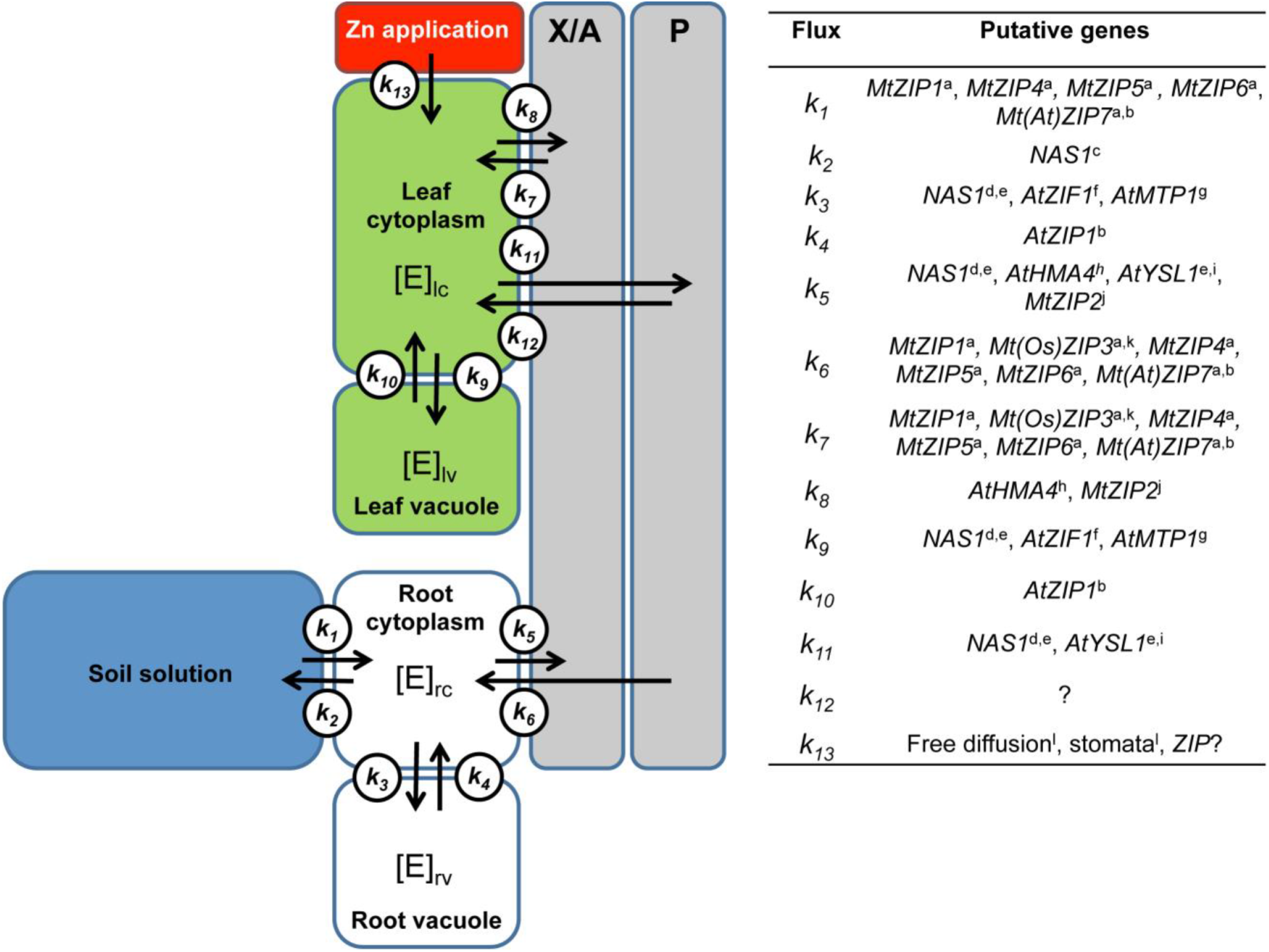
Suggested model for the roles of putative genes encoding proteins involved in Zinc (Zn) transport-related processes in alfalfa (*Medicago sativa*). The sites of action in the plant (i.e., root cytoplasm, rc; root vacuole, rv; xylem and apoplast, X/A; phoelm, P; leaf cytoplasm, lc; leaf vacuole, lv) and the element (E) fluxes (K_1-13_) are reported. The concentration of the element is indicated in each site [E]. The scheme synthetizes information across studies in various plants. Gene abbreviations: *ZIP*, Zrt-/Irt-like Protein; *NAS*, Nicotianamine synthase; *ZIF*, Zinc-Induced Facilitator; *MTP,* Metal Transporter Protein; *HMA,* P_1B_-type Heavy Metal ATPase; *YSL*, Yellow Stripe Like Protein; *ZIP*? indicates a generic *ZIP*; free diffusion: diffusion through leaf epidermis; stomata: absorption through stomata. Plant abbreviations: Mt, *Medicago truncatula*; At*, Arabidopsis thaliana;* Os, *Oryza sativa*. References: ^a^López-Millán et al., 2004; ^b^Milner et al., 2013; ^c^Aarts, 2014; ^d^Clemens et al., 2013; ^e^Curie et al., 2009; ^f^Haydon et al., 2012; ^g^Desbrosses-Fonrouge et al., 2005; ^h^Hussain et al., 2004; ^i^Palmer and Guerinot, 2009; ^j^Burleigh et al., 2003; ^k^Sasaki et al.,? 2015; ^l^lFageria et al., 2009.

Using the draft genome sequence of alfalfa in the Alfalfa Gene Index and Expression Atlas Database (AEGD) (O’Rourke et al., 2015; http://plantgrn.noble.org/AGED/index.jsp), homologous gene sequences of *M. sativa* were retrieved by BLASTn similarity searches using the gene sequences of *M. truncatula*. The chosen genes for *M. sativa* were named *MsZIP_1-7_* for the seven *ZIP* genes and *MsNAS1*, *MsHMA4*, *MsZIF1*, *MsYSL1*, *MsMTP1* for the other selected genes. The two reference genes were named *MsACT-101* and *MsEF1-α*. The gene sequences and their annotations have been deposited in NCBI under the Submission # 2338923.

Forward and reverse new PCR primers for the 12 Zn transport-related genes and the two reference genes suitable for SYBR^®^ Green II RT-qPCR assays (Biorad, USA) were designed (Table 2). The Primer-BLAST online tool in the National Center for Biotechnology Information (NCBI; https://www.ncbi.nlm.nih.gov/tools/primer-blast/) was used to design primers. The newly designed RT-qPCR assays are suitable for both *M. sativa* and *M. truncatula*. The length of the fragment, the Sanger sequences of the PCR amplicons (Table 2) and the single melting temperature peaks confirmed the specificity of the new RT-qPCR assays (Supplemental Fig. S4). Sanger sequencing was performed on PCR amplicons of three cDNA samples (Supplemental Material and Methods S1). Examples of electropherograms of the sequences are reported in Supplemental Fig. S5. The sequences of the obtained PCR amplicons have been deposited in NCBI the Submission # 2338930. Amplification efficiencies (E) in the range of 96.1-111.0% are evidence of accurate quantification, while the coefficients of correlation (R^2^>0.998) indicate high precision of measurements across concentration ranges of at least 3-4 orders of magnitude (Table 2; Supplemental Fig. S6). The concentration ranges over which the relationship between the relative fluorescence and the logarithm of the concentration is linear, and the precision of quantification (standard curves) as reflected in the coefficient of correlation (R^2^), were determined using three independent 10-fold serial dilutions of a cDNA sample of *M. sativa*. The accuracy of quantification was determined by the efficiency (E) of each qPCR amplification, using the equation *E* = [10^-1/S^ − 1] × 100, where S is the slope of the standard curve. The evaluation of the reference genes based on the cycle threshold (Ct) values made us choose the actin gene (*MsACT-101*) for quantifying relative gene expression in the shoots and the elongation factor 1-α (*MsEF1-α*) gene for quantifying relative gene expression in roots (Supplemental Fig. S7a,b). This choice was based on the observations that there was no statistical difference in the expression of the reference genes in tissues following foliar Zn applications and that *MsACT-101* and *MsEF1-α* showed the smallest overall variation in the shoot and root, respectively (Supplemental Fig. S7c,d).

**Table 2.**
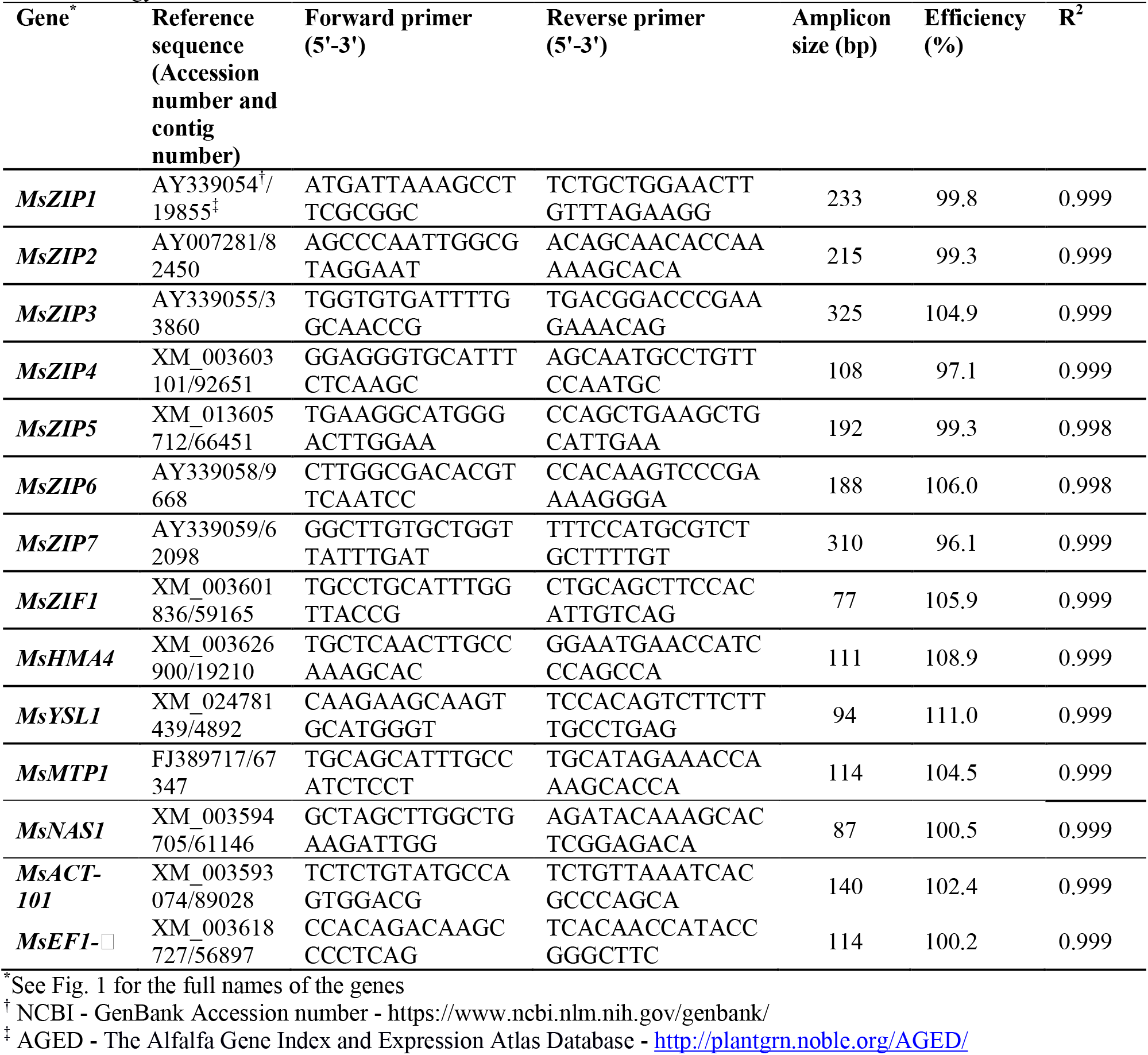
Gene name, forward and reverse sequences of fourteen newly designed primer pairs for the quantification of the expression of genes of alfalfa (*Medicago sativa),* encoding proteins involved in cellular zinc (Zn) influx and efflux and Zn chelation (see **Fig. 1**). Two reference genes (i.e., *MsACT-101* and *MsEF1*-□□□ were also designed. The length of the amplicons, the primer amplification Efficiency (%) and R^2^ of the standard curve are indicated (see **Fig. S3**). The reference sequences are indicated by the accession number of the *Medicago truncatula* sequences and by the contig number of the *M. sativa* sequences. The primers were designed using the National Center for Biotechnology Information *PRIMER Blast* online tool.

### RNA extraction and gene expression analysis

Total RNA was extracted from 50 mg subsamples of fresh shoot and root tissue, using the RNeasy Mini Kit (Qiagen, Hilden, Germany). The extractions were performed from tissues of plants treated with the foliar Zn doses that produced a significant increase in Zn concentration in shoots (0.1, 1 and 10 mg Zn plant^-1^) and the control plants to which no foliar Zn had been applied (24 RNA extractions). Any DNA in the RNA extracts was removed by a DNase treatment (Promega, USA). The purity of the RNA extracts was verified by spectroscopic light absorbance measurements at 230 nm, 260 nm and 280 nm using the NanoDrop 2000 (Termo Scientific, Worchesre, MA, USA) (Desjardins and Conkin, 2010). The integrity and approximate concentration of the extracted RNA was determined by electrophoresis of the RNA extracts in a 1% agarose gel containing Sybr Safe (Invitrogen, Carlsbad, CA). One microgram of total RNA was reverse transcribed to complementary DNA (cDNA) using the iScript cDNA Synthesis Kit (Biorad, Hercules, California) in a 20 μL reaction volume. The RT-qPCRs for gene expression analysis were run as three technical replicates with a final reaction volume of 20 μL, containing 10 μL of SYBR Green Supermix (Biorad), 5 μL of 100-fold diluted cDNA, and 0.4 μM final concentrations of the gene-specific PCR primers on a CFX Connect Real-Time System thermal cycler (Biorad, Hercules, California). The qPCR conditions were 95°C for 3’, followed by 40 cycles of 95° C for 5’, and 60° C for 30’’. A dissociation curve of each reaction was performed (65° C to 95° C, 0.5° C increment every 5’’) to check that PCR amplified only one product. The most suitable reference gene for relative gene expression analysis was determined by comparing the expression levels of the reference genes *MsACT-101* and *MsEF1-α* across all cDNA samples. Relative gene expression was calculated using the double standardisation (ΔΔCq) method that requires a reference gene and a control treatment (Livak and Schmittgen, 2001).

### Bioinformatic and statistical analyses

A BLAST search was performed in the Alfalfa Gene Index and Expression Atlas database using the *ZIP1-7, ZIF1, MTP1, YSL1, HMA4* and *NAS* coding sequences from *M. truncatula*. This allowed the identification of gene sequences encoding potential metal transporters and chelators in the whole *M. sativa* genome. The sequences obtained were aligned with the corresponding sequences from *M. truncatula* and the length of the *M. sativa* genes were determined after removing the external unaligned nucleotides. The *M. sativa* and *M. truncatula ZIP* gene sequences were also aligned with those of other plant species (*A. thaliana, G. max, H. vulgare, O. sativa, Triticum aestivum,* and *Zea mays*) obtained from a search of GenBank. Similarly, the *M. sativa* and *M. truncatula* gene sequences of *ZIF1, MTP1, YSL1, HMA4* and *NAS* were aligned with their corresponding sequences of other plant species (*A. thaliana, G. max, H. vulgare, O. sativa, T. aestivum,* and *Z. mays*) obtained from a search of GenBank. Sequence alignments were performed using the algorithm ClustalW in MEGA X (Kumar et al., 2018). Phylogenetic comparisons were performed to infer the putative roles of the selected *M. sativa* Zn transport-related proteins. The phylogenetic trees were inferred by Neighbor-Joining (NJ) analysis (Saitou et al., 1987) in MEGA X and the evolutionary distances were calculated using the p-distance method (Nei and Kumar, 2000). Branch support bootstrap values were derived from 500 bootstrap replicates. The phylograms were drawn by MEGA X and edited using Adobe Illustrator CC 2017.

The effect of the application of the foliar Zn on tissue Zn concentration and on the expression of the selected genes was analysed in shoots and roots separately by one-way analysis of variance (ANOVA), followed by a Tukey-B test in the case of significance of the response to foliar Zn application. When required, gene expression data were log-transformed to meet the ANOVA assumptions. The data displayed graphically are the means and associated standard errors of the untransformed raw data. All statistical analyses were performed using the software package SPSS version 21.0 (SPSS Inc., Chicago, IL, USA). Permutational analysis of variance (PERMANOVA; Anderson, 2001) was used to test the effect of foliar Zn application and plant organ (shoot and root) on the expression of the seven *ZIP* genes and of the other five genes encoding Zn transport-related processes separately. In addition, the PERMANOVA was performed on the expression of all the genes together. The response data matrices were standardised by sample, and total and then Euclidean distances were calculated among samples. *P*-values were calculated using the MonteCarlo test (Anderson and Braak, 2003). Since PERMANOVA is sensitive to differences in multivariate location and dispersion, analysis of homogeneity of multivariate dispersion (PERMDISP; Anderson, 2006) was performed to check the homogeneity of dispersion among groups. The analyses were performed using PRIMER 7 and PERMANOVA+ software (Clarke and Gorley, 2015). Finally, heatmaps were constructed to illustrate correlations in expression among *ZIP*s and among other genes encoding Zn transport-related processes using the R package ggplot2 (Wickham, 2011), using the average linkage clustering of the Pearson correlations calculated from relative gene expression following foliar Zn application.

## Supplemental Materials

**Supplemental Figure S1.** Shoot and root zinc (Zn) content of alfalfa (*Medicago sativa* L.) five days after application of six doses of Zn to leaves.

**Supplemental Figure S2.** Neighbor-Joining phylogenetic tree of *ZIP* gene sequences of *Medicago sativa*, *Medicago truncatula* and other plant species.

**Supplemental Figure S3.** Neighbor-Joining phylogenetic trees of sequences of Zinc Induced Facilitator (*ZIF),* Metal Tollerance Protein (*MTP*), Yellow Stripe Like protein (*YSL*), Heavy Metal Transporter (*HMA*) and Nicotianamine Synthase (*NAS*) genes of *Medicago sativa, Medicago truncatula* and other plant species.

**Supplemental Figure S4.** Melting curve analysis of the qPCR products obtained by the newly designed pair of primers.

**Supplemental Figure S5.** Electropherograms of PCR products obtained by the newly designed primers for *MsZIP_1-7_* transporters and for other Zn related genes.

**Supplemental Figure S6.** Standard curves for the newly designed qPCR primer pairs.

**Supplemental Figure S7.** Cycle threshold value of the reference genes, *actin 101 (MsACT-101*) and *elongation factor 1-α (MsEFf1-α*) in shoots and roots for the no-Zn addition control and the three Zn doses.

**Supplemental Material and Methods S1.** PCR amplification conditions.

## ACKNOWLEDGEMENTS

We acknowledge the Italian Institute of Technology (IIT) for funding for PhD Fellowship of AC under the PhD programme in Agrobiosciences at the Scuola Superiore Sant’Anna of Pisa, Italy. PJW was supported by the Rural and Environment Science and Analytical Services Division (RESAS) of the Scottish Government. We acknowledge Dr. Hannes A. Gamper for the technical support in real-time RT-PCR.

